# Outpatient prescribing of four major antibiotic classes and prevalence of antimicrobial resistance in US adults

**DOI:** 10.1101/456244

**Authors:** Edward Goldstein, Derek R. MacFadden, Robyn S. Lee, Marc Lipsitch

## Abstract

**Background:** There is limited information on the relation between antibiotic use and antibiotic resistance in the US.

**Methods:** We used multivariable logistic regression to relate state-specific rates of outpatient prescribing overall for fluoroquinolones, penicillins, macrolides, and cephalosporins between 2011-2014 to state-specific prevalence of resistance for select combinations of antibiotics/bacteria among catheter-associated urinary tract infections (CAUTIs) in the CDC Patient Safety Atlas data between 2011-2014 for adults aged 65+y and 19-64y, adjusting for potential confounders.

**Results:** Rates of fluoroquinolone prescribing were positively associated with prevalence of fluoroquinolone resistance in *Escherichia coli* and *Pseudomonas aeruginosa* (both age groups), resistance to extended-spectrum (ES) cephalosporins in *E. coli* (aged 19-64y), and resistance to methicillin in *Staphylococcus aureus* (aged 19-64y). Rates of penicillin prescribing were positively associated with prevalence of resistance to fluoroquinolones in *E. coli* (aged 65+) and *P. aeruginosa* (both age groups), and resistance to ES cephalosporins in *Klebsiella* spp. (both age groups). Rates of cephalosporin prescribing were negatively associated with prevalence of resistance to fluoroquinolones in *E. coli* and resistance to ES cephalosporins in *Klebsiella* spp. (both age groups). Average annual temperature was positively associated with prevalence of resistance to ES cephalosporins in *E. coli* and *P. aeruginosa,* and resistance to fluoroquinolones in *E. coli.*

**Conclusions:** Our results suggest that prescribing of fluoroquinolones and penicillins to US adults is associated with prevalence of antibiotic resistance, including ESBLs and MRSA. Further work is needed to understand the potential benefit of replacing fluoroquinolones and penicillins by other antibiotics for reducing prevalence of antibiotic resistance.

## Introduction

Antibiotic resistance is a major public health threat that is receiving increasing attention and in some areas is growing rapidly [1], with antibiotic resistance and use contributing to a variety of illness outcomes, including sepsis [2,3]. Several studies have suggested that levels of antibiotic prescribing affect the prevalence of antibiotic resistance [4-10]. At the same time, the effects of antibiotic prescribing on antibiotic resistance are non-uniform, and may depend on the antibiotic/bacteria combination and other factors, e.g. [4,8,11]. Antibiotic prescribing may also affect the prevalence of bacterial phenotypes resistant to other antibiotics. For example, fluoroquinolone use was found to be positively associated with prevalence of MRSA infections [12-15], while amoxicillin use was positively associated with prevalence of trimethoprim resistance in Enterobacteriaceae in England [5]. High prevalence of resistance to commonly prescribed antibiotics, particularly fluoroquinolones and penicillins in certain bacteria (*E. coli, P. aeruginosa, Klebsiella* spp.), as well as of multidrug resistance and cross-resistance with ESBLs is documented in various studies, e.g. [16-19,10]. It is therefore conceivable that use of commonly prescribed antibiotics in the US such as fluoroquinolones and penicillins could contribute to the prevalence of multidrug resistance and ESBLs in certain bacteria, particularly those for which prevalence of fluoroquinolone and penicillin resistance is high (e.g. Table 2 in [16]). For example, a US study of *E. coli*-related UTIs in hospitalized patients [17] suggests a prevalence of resistance to fluoroquinolones of 34.5%, prevalence of resistance to 3^rd^ generation cephalosporins (which generally serves as a marker for ESBL) of 8.6%, and prevalence of joint resistance of 7.3%. Thus it is plausible that fluoroquinolone use contributes to prevalence of resistance to 3^rd^ generation cephalosporins in *E. coli*-related UTIs in the US, which may be similar to the effect of amoxicillin use on trimethoprim resistance in Enterobacteriaceae in England [5]. We note that evidence in the literature about the relation between antibiotic use in the US and antibiotic resistance, particularly use of one antibiotic and resistance to another, is still limited. Further understanding of the relation between antibiotic consumption, resistance, and the associated severe bacterial infections, particularly in the US context, is needed to inform antibiotic prescribing guidelines and practices. Such understanding is especially important in light of the high volume of inappropriate antibiotic prescribing in the US [20- 22], including for fluoroquinolones [23].

Previous work suggests that outpatient prescribing of antibiotics plays a key role in the propagation of antibiotic resistance [24]. In this paper, we examine the relationship between outpatient prescribing for different antibiotic classes and prevalence of antibiotic resistance for certain combinations of antibiotics/bacteria in the US. We use a regression framework to relate the state-specific rates of outpatient prescribing of fluoroquinolones, penicillins, macrolides, and cephalosporins recorded in the US CDC Healthcare Safety Network (NHSN) Antibiotic Resistance Patient Safety Atlas prescribing data [25], and state-specific prevalence of antibiotic resistance for different combinations of antibiotics/bacteria in hospitalized elderly and non-elderly adults documented in the US CDC Antibiotic Resistance Patient Safety Atlas resistance data [26,27]. In particular, the aim is to examine the relation between rates of prescribing for commonly used antibiotics and prevalence of resistance to fluoroquinolones, Extended Spectrum (ES) cephalosporins (which generally serves as a marker for ESBL), and of resistance to methicillin in *S. aureus*, as suggested in the previous paragraph.

## Methods

### Data

We extracted data on the annual state-specific per capita rates of outpatient prescribing for four classes of antibiotics: fluoroquinolones, penicillins, macrolides, and cephalosporins between 2011 and 2014 from the US CDC Antibiotic Resistance Patient Safety Atlas [25]. Average annual state-specific outpatient prescribing rates (per 1,000 residents) for each class of antibiotics between 2011-2014 were then estimated as the average for the corresponding annual rates.

Annual state-specific population estimates in each age group of adults were obtained as the yearly July 1 population estimates in [28]. State-specific population in each age group between 2011-2014 was then estimated by averaging the corresponding annual population estimates. Average population densities for the different states between 2011-2014 were obtained by dividing the average annual population estimates between 2011-2014 by the state land area [29]. Data on median household income for the different US states between 2011-2014 were extracted from [30]. Data on average monthly temperature for the different US states were obtained from [31].

We extracted data on the prevalence of antibiotic resistance for bacterial specimens collected from hospitalized patients in the US between 2011-2014 from the US CDC National Healthcare Safety Network (NHSN) Antibiotic Resistance Patient Safety Atlas [26,27]. Those data are stratified by age group (<1y,1-18y,19-64y, 65+y), state, year, infection type (CAUTI, CLABSI, SSI, [26]), and combination of bacteria/antibiotics (31 combinations documented in [26]). The following seven combinations of antibiotics/bacteria were studied in the paper: resistance to fluoroquinolones in *E. coli* and *P. aeruginosa*, resistance to ES cephalosporins in *E. coli*, *P. aeruginosa*, *Klebsiella* spp. and *Enterobacter*, and resistance to methicillin in *S. aureus*.

Resistance prevalence varies by type of infection (Tables 6, 7, 9 in [27]), making the combination of different types of infection into the analysis problematic. Moreover, for a given combination of bacteria/antibiotics, and a given type of infection, only a fraction of states reported the corresponding resistance pattern [26], with that fraction generally being highest for catheter-associated urinary tract infections (CAUTIs). Therefore, for each combination of bacteria/antibiotics and age group of adults (19-64y or 65+y), data on resistance in only the CAUTI samples for the given age group/combination of bacteria/antibiotics in [26] for those states that reported the corresponding aggregate data between 2011-2014 were included in the analysis.

### Statistical analysis

There is evidence in the literature that prevalence of antibiotic resistance is affected not only by the levels of antibiotic prescribing but also by a variety of additional factors, such as socio-demographic factors influencing bacterial transmission, both in the community and during the course of medical treatment, and temperature that is related to the rate of bacterial growth [11,32]. Correspondingly, we have adjusted the analysis of the relation between the rates of antibiotic prescribing and prevalence of antibiotic resistance by including additional covariates described below.

For each of the two age groups of adults (19-64y and 65+y) and each of the seven combinations of antibiotics/bacteria described in the previous subsection (e.g. resistance to fluoroquinolones in *E. coli*), we used a multivariable logistic regression model with random effects to relate the state-specific outpatient prescribing rates for fluoroquinolones, penicillins, macrolides, and cephalosporins between 2011-2014 [25] to the state-specific prevalence of resistance between 2011-2014 for the given combination of antibiotics/bacteria in CAUTI samples from the given age group [26], adjusting for median household income, average annual temperature, and population density as additional predictors of resistance. Specifically, for each state *s* that reported the corresponding aggregate resistance data between 2011-2014 in [26], for each tested CAUTI sample from that state, the binary outcome is the presence of resistance, and the covariates are: *A_i_*(*S*) (*i* = 1,…,4), the state-specific average annual outpatient prescribing rates between 2011-2014, per 10 state residents, for the four studied classes of antibiotics; *I*(*s*), the median state-specific household income between 2011-2014; *T*(*s*), the average annual temperature (°F) between 2000-2014 for the state; *PD*(*s*), the state-specific average population density between 2011-2014. Additionally, we include random effects *RE* for the ten Health and Human Services (HHS) regions in the US. Those effects partly adjust for the correlation between the prescribing rates for different antibiotics at the regional level, as well as for additional factors potentially not accounted for by the predictors used in the model, such as demographic factors and possible geographic differences in testing practices for resistance in CAUTI samples [26]. Thus we model the probability of resistance for a CAUTI sample from state *s* as

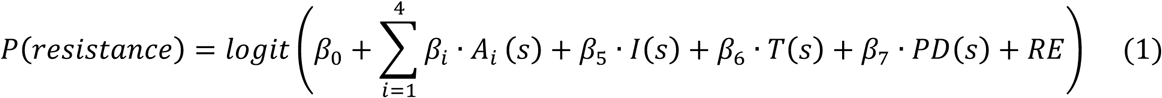

## Results

Table 1 shows the estimates (median and the lower/upper quartiles for the states reporting the corresponding data) for (i) the number of CAUTI samples tested; (ii) the number of resistant CAUTI samples; and (iii) the prevalence of resistance for CAUTI samples collected between 2011-2014 ([26]) for each of the seven combinations of bacteria/antibiotics considered in this paper for adults aged 19-64y and 65+y. Table 2 shows the correlations between the state-specific rates of outpatient prescribing for each of the four major antibiotics classes (fluoroquinolones, penicillins, cephalosporins and macrolides) between 2011-2014 [25] and the state-specific prevalence of resistance for each of the seven combinations of bacteria/antibiotics considered in this paper for adults aged 19-64y and 65+y for the states reporting the corresponding data on CAUTI samples in [26]. We note the large number of positive correlations that may partly reflect the strong association between rates of prescribing for the different pairs of antibiotics [3].

**Table 1:** Estimates (median and the lower/upper quartiles) for (i) the number of tested CAUTI samples; (ii) the number of resistant CAUTI samples; and (iii) the prevalence of resistance for CAUTI samples collected between 2011-2014 for the seven combinations of bacteria/antibiotics considered in this paper for adults aged 19-64y and 65+y for the states reporting the corresponding data on CAUTI samples in [26].

|  | Aged 19-64y |  |  | Aged 65+y |  |  |
| --- | --- | --- | --- | --- | --- | --- |
|  | Tested samples | Resistant samples | Resistance prevalence | Tested samples | Resistant samples | Resistance prevalence |
| *E. coli* / fluoroquinolones | 232.5<br>[79.2,374.8] | 73<br>[24.2,118.2] | 30.3%<br>[25.6%,35.7%] | 238<br>[90.5,441] | 80<br>[21.5,141.5] | 32.3%<br>[23.6%,36.8%] |
| *E. coli* / ES cephalosporins | 197<br>[80,330] | 21<br>[9,43] | 10.1%<br>[7.6%,14.1%] | 210<br>[108,364] | 25<br>[8,47] | 11.3%<br>[8.6%,13.8%] |
| *P. aeruginosa* / fluoroquinolones | 104<br>[55.5,200.5] | 34<br>[18.5,61] | 31.4%<br>[24.1%,35.2%] | 117<br>[59,237] | 28<br>[11,58] | 23.3%<br>[17.8%,28.2%] |
| *P. aeruginosa* / ES cephalosporins | 105<br>[54,200] | 16<br>[7,25] | 12.4%<br>[9.8%,14.8%] | 116<br>[58,237] | 12<br>[5,28] | 9.6%<br>[7.7%,12.7%] |
| *Klebsiella spp.* / ES cephalosporins | 84<br>[41,167] | 15<br>[5,44.5] | 15.9%<br>[11.1%,23.4%] | 109<br>[48,176] | 13.5<br>[3.8,42.2] | 12.6%<br>[6.6%,19.1%] |
| Enterobacter / ES cephalosporins | 54.5<br>[33.2,79.2] | 21.5<br>[11,30.2] | 32.9%<br>[27.1%,38.7%] | 47<br>[33,76] | 16<br>[9,27.5] | 32.4%<br>[28.8%,36.6%] |
| *S. aureus* / methicillin | 29.5<br>[23,47] | 14<br>[11,26.5] | 45.1%<br>[41.6%,51.6%] | 33<br>[27.8,43.8] | 21.5<br>[18,27.5] | 61.6%<br>[55.2%,66.7%] |

**Table 2:** Correlations (with 95% CIs) between the state-specific rates of outpatient prescribing for fluoroquinolones, penicillins, cephalosporins and macrolides between 2011- 2014 [25] and the state-specific prevalence of resistance for the seven combinations of bacteria/antibiotics considered in this paper for adults aged 19-64y and 65+y for the states reporting the corresponding data on CAUTI samples in [26].

|  | *E. coli* R to<br>fluoroquinolones | *E. coli* R to<br>ES cephalosporins | *P. aeruginosa* R to<br>fluoroquinolones | *P. aeruginosa* R to<br>ES cephalosporins | *Klebsiella* spp. R to<br>ES cephalosporins | Enterobacter R to<br>ES cephalosporins | Methicillin R<br>among *S. aureus* |
| --- | --- | --- | --- | --- | --- | --- | --- |
|  | Aged 19-64y |  |  |  |  |  |  |
| Fluoroquinolones | 0.68<br>(0.5,0.81) | 0.14<br>(-0.14,0.41) | 0.51<br>(0.23,0.71) | 0.42<br>(0.13,0.65) | 0.16<br>(-0.15,0.44) | 0.22<br>(-0.14,0.53) | 0.36<br>(-0.13,0.71) |
| Penicillins | 0.48<br>(0.23,0.67) | 0.07<br>(-0.22,0.34) | 0.41<br>(0.11,0.64) | 0.25<br>(-0.06,0.52) | 0.03<br>(-0.28,0.33) | 0.14<br>(-0.22,0.46) | -0.17<br>(-0.59,0.32) |
| Cephalosporins | 0.43<br>(0.17,0.63) | -0.12<br>(-0.39,0.17) | 0.39<br>(0.09,0.63) | 0.22<br>(-0.09,0.5) | -0.19<br>(-0.47,0.12) | 0.06<br>(-0.3,0.4) | 0.09<br>(-0.4,0.53) |
| Macrolides | 0.61<br>(0.39,0.76) | 0.05<br>(-0.24,0.33) | 0.44<br>(0.14,0.66) | 0.34<br>(0.03,0.58) | 0.06<br>(-0.25,0.35) | 0.14<br>(-0.22,0.47) | 0.09<br>(-0.39,0.54) |
|  | Aged 65+y |  |  |  |  |  |  |
| Fluoroquinolones | 0.59<br>(0.37,0.74) | 0.17<br>(-0.12,0.43) | 0.54<br>(0.29,0.72) | 0.09<br>(-0.21,0.38) | 0.19<br>(-0.11,0.46) | 0.5<br>(0.2,0.72) | 0.37<br>(-0.04,0.67) |
| Penicillins | 0.44<br>(0.18,0.64) | 0.09<br>(-0.19,0.37) | 0.43<br>(0.15,0.64) | 0.06<br>(-0.24,0.35) | 0.11<br>(-0.19,0.39) | 0.27<br>(-0.07,0.55) | 0.19<br>(-0.23,0.55) |
| Cephalosporins | 0.34<br>(0.07,0.56) | -0.07<br>(-0.34,0.22) | 0.27<br>(-0.02,0.53) | 0.07<br>(-0.23,0.35) | -0.08<br>(-0.37,0.22) | 0.22<br>(-0.12,0.52) | 0.27<br>(-0.15,0.61) |
| Macrolides | 0.47<br>(0.22,0.66) | 0.09<br>(-0.19,0.37) | 0.41<br>(0.13,0.63) | 0.08<br>(-0.22,0.37) | 0.12<br>(-0.19,0.4) | 0.51<br>(0.21,0.72) | 0.25<br>(-0.18,0.59) |

Estimates of the coefficients of the regression model given by eq. 1 for the different age groups and combination of antibiotics/bacteria are presented in Table 3 (aged 65+y) and Table 4 (aged 19-64y). Tables 3 and 4 show that rates of fluoroquinolone prescribing were positively associated with prevalence of fluoroquinolone resistance in *E. coli* (both age groups) and *P. aeruginosa* (both age groups), prevalence of resistance to ES cephalosporins in *E. coli* (aged 19-64y), and prevalence of resistance to methicillin in *S. aureus* (aged 19- 64y). Rates of penicillin prescribing were positively associated with prevalence of resistance to fluoroquinolones in *E. coli* (aged 65+) and *P. aeruginosa* (both age groups), and prevalence of resistance to ES cephalosporins in *Klebsiella* spp. (both age groups). Rates of cephalosporin prescribing were negatively associated with prevalence of fluoroquinolone resistance in *P. aeruginosa*, and prevalence of resistance to ES cephalosporins in *Klebsiella* spp. (both age groups). Additionally, average annual temperature was positively associated with prevalence of resistance to ES cephalosporins in *E. coli* (aged 65+y) and *P. aeruginosa* (both age groups), and prevalence of resistance to fluoroquinolones in *E. coli* (aged 65+y).

**Table 3:** Estimates of the regression coefficients (multiplied by 100 for temperature) in the model given by eq. 1 for resistance for the different combinations of antibiotics/bacteria in adults aged 65+y. The regression coefficients for the antibiotic prescribing rates indicate the logarithm of the odds ratio for resistance when the rate of prescribing of a given antibiotic (per 10 individuals) increases by 1.

|  | *E. coli* R to<br>fluoroquinolones | *E. coli* R to<br>ES cephalosporins | *P. aeruginosa* R to<br>fluoroquinolones | *P. aeruginosa* R to<br>ES cephalosporins | *Klebsiella* spp. R to<br>ES cephalosporins | Enterobacter R to<br>ES cephalosporins | Methicillin R among<br>*S. aureus* |
| --- | --- | --- | --- | --- | --- | --- | --- |
| Fluoroquinolones<br>(prescription per 10<br>residents/y) | **0.76**<br>**(0.08,1.4)** | 0.84<br>(-0.18,1.9) | **1**<br>**(0.02,2)** | -0.21<br>(-1.7,1.2) | 0.02<br>(-1.7,1.7) | 0<br>(-1.2,1.2) | 1.7<br>(-0.34,3.7) |
| Penicillins<br>(prescription per 10<br>residents/y) | **0.32**<br>**(0.01,0.63)** | -0.04<br>(-0.53,0.44) | **0.88**<br>**(0.38,1.4)** | 0.67<br>(-0.03,1.4) | **1.7**<br>**(0.83,2.5)** | -0.26<br>(-1.1,0.56) | 0.71<br>(-0.47,1.9) |
| Cephalosporins<br>(prescription per 10<br>residents/y) | -0.24<br>(-0.52,0.03) | -0.42<br>(-0.83,0) | **-0.65**<br>**(-1.1,-0.22)** | 0.11<br>(-0.47,0.69) | **-2**<br>**(-2.6,-1.3)** | -0.29<br>(-1,0.45) | -0.52<br>(-1.7,0.64) |
| Macrolides<br>(prescription per 10<br>residents/y) | -0.23<br>(-0.58,0.12) | -0.09<br>(-0.64,0.47) | -0.52<br>(-1.1,0.06) | -0.59<br>(-1.4,0.23) | -0.15<br>(-1,0.71) | 0.63<br>(-0.26,1.5) | -0.77<br>(-2.4,0.84) |
| Median household income (\$100,000) | 0.62<br>(-0.08,1.3) | **1.5**<br>**(0.41,2.5)** | 0.09<br>(-1,1.2) | 0.67<br>(-0.91,2.3) | -2.4<br>(-4.9,0.17) | **-0.47**<br>**(-0.72,-0.22)** | **1.4**<br>**(1,1.7)** |
| Average annual temperature (°F) | **2.4**<br>**(1.6,3.3)** | **1.3**<br>**(0.13,2.5)** | 0.97<br>(-0.22,2.2) | **1.9**<br>**(0.18,3.6)** | 2.4<br>(-0.15,5) | 0.93<br>(-0.51,2.4) | 2.1<br>(-0.27,4.4) |
| Population density (10,000 inhabitants/sq. mile) | -0.57<br>(-1.3,0.12) | -0.45<br>(-1.5,0.55) | -0.21<br>(-1.1,0.72) | 0.39<br>(-0.93,1.7) | 0.57<br>(-0.63,1.8) | 4.4<br>(-0.44,9.3) | -5.9<br>(-13.4,1.5) |

**Table 4:** Estimates of the regression (multiplied by 100 for temperature) in the model given by eq. 1 for resistance for the different combinations of antibiotics/bacteria in adults aged 19-64y. The regression coefficients for the antibiotic prescribing rates indicate the logarithm of the odds ratio for resistance when the rate of prescribing of a given antibiotic (per 10 individuals) increases by 1.

|  | E. coli R to fluoroquinolones | E. coli resistant to ES cephalosporins | P. aeruginosa R to fluoroquinolones | P. aeruginosa R to ES cephalosporins | Klebsiella spp. R to ES cephalosporins | Enterobacter R to ES cephalosporins | Methicillin R among *S. aureus* |
| --- | --- | --- | --- | --- | --- | --- | --- |
| Fluoroquinolones (prescription per 10 residents/y) | **1.4**<br>**(0.59,2.1)** | **1.5**<br>**(0.23,2.7)** | **1.2**<br>**(0.17,2.2)** | -0.32<br>(-1.3,0.67) | 1<br>(-0.63,2.6) | -0.07<br>(-1.3,1.2) | **2.8**<br>**(0.66,4.9)** |
| Penicillins (prescription per 10 residents/y) | -0.24<br>(-0.58,0.09) | -0.07<br>(-0.6,0.46) | **0.89**<br>**(0.36,1.4)** | 0.02<br>(-0.68,0.71) | **0.82**<br>**(0.07,1.6)** | 0.01<br>(-0.79,0.81) | -0.97<br>(-2.4,0.45) |
| Cephalosporins (prescription per 10 residents/y) | -0.24<br>(-0.54,0.07) | -0.64<br>(-1.1,-0.18) | **-0.6**<br>**(-1.1,-0.15)** | 0.29<br>(-0.27,0.86) | **-1.7**<br>**(-2.4,-1.1)** | 0.72<br>(-0.09,1.5) | -0.18<br>(-1.7,1.3) |
| Macrolides (prescription per 10 residents/y) | -0.09<br>(-0.48,0.31) | -0.09<br>(-0.72,0.55) | -0.61<br>(-1.2,0.01) | 0.09<br>(-0.66,0.84) | 0.06<br>(-0.83,0.95) | -0.56<br>(-1.5,0.38) | -0.18<br>(-1.7,1.3) |
| Median household income (\$100,000) | 0.37<br>(-0.41,1.1) | **1.9**<br>**(0.74,3.1)** | -0.11<br>(-1.3,1.1) | **0.52**<br>**(0.27,0.77)** | 0.15<br>(-1.4,1.7) | **-2.3**<br>**(-2.6,-2.1)** | **4.1**<br>**(3.6,4.5)** |
| Average annual temperature (°F) | 0.8<br>(-0.1,1.7) | 0.26<br>(-1.1,1.6) | 0.18<br>(-1,1.4) | **2.2**<br>**(0.9,3.5)** | -0.08<br>(-2,1.8) | -0.62<br>(-2.2,0.96) | 0.67<br>(-2,3.3) |
| Population density (10,000 inhabitants/sq. mile) | -0.26<br>(-0.92,0.41) | 0.11<br>(-0.78,1) | -0.61<br>(-1.5,0.31) | 0.61<br>(-0.42,1.6) | 0.67<br>(-0.48,1.8) | **13.4**<br>**(8.1,18.6)** | -1.4<br>(-15.6,12.8) |

## Discussion

There is mixed evidence in the literature about the relationship between antibiotic prescribing and antibiotic resistance. A number of studies have shown a relation between rates of antibiotic prescribing and prevalence of antibiotic resistance [4-10], including resistance to other antibiotics [5,12-15]. At the same time, the strength of the relation between antibiotic prescribing and prevalence of resistance is likely to be non-uniform and may depend on the pairing of bacteria/antibiotic and other factors [4,8,11]. Better understanding of these relationships in the US context should help inform antibiotic prescribing guidelines and improve antibiotic prescribing practices. Such understanding is particularly important in light of both the relation between the use of/resistance to different antibiotics, including penicillins and fluoroquinolones and rates of severe bacterial infections, including those resulting in sepsis [2,3], as well as the high volume of inappropriate antibiotic prescribing in the US [20-23]. In this study, multivariable logistic mixed-effects regression is used to evaluate the association between the levels of outpatient prescribing for four major classes of antibiotics (fluoroquinolones, penicillins, cephalosporins, and macrolides) and prevalence of resistance for certain combinations of antibiotics/bacteria. Our results suggest that rates of outpatient use of commonly prescribed antibiotics, particularly penicillins and fluoroquinolones, are associated not only with prevalence of fluoroquinolone resistance, but also with prevalence of resistance to ES cephalosporins (which generally serves as a marker for ESBL), and prevalence of MRSA. Those results resonate with our earlier findings about the associations between the rates of prescribing of penicillins and fluoroquinolones and rates of septicemia hospitalization in US adults aged over 50y [3]. Along with evidence of high levels of inappropriate antibiotic prescribing [20-23], this strongly supports the need to change the practices related to antibiotic prescribing, particularly for penicillins and fluoroquinolones in the US. Our results also support earlier findings [32] about the relation between temperature and antibiotic resistance prevalence, with positive associations found in this study between average annual temperature and prevalence of antibiotic resistance in certain bacteria (*E. coli* and *P. aeruginosa*).

In our previous work [3], we identified associations between the rates of outpatient prescribing overall for certain classes of antibiotics, particularly penicillins and fluoroquinolones, and rates of septicemia hospitalization in US adults aged over 50y. Such associations may be mediated by antibiotic resistance through direct or indirect mechanisms. Specifically, bacterial infections resistant to a given antibiotic may fail to clear following treatment with that antibiotic, eventually devolving into sepsis. Additionally, use of a given antibiotic may contribute to the prevalence of resistance to another antibiotic, potentially also leading to poor outcomes following treatment with the latter drug. For example, amoxicillin use was found to be associated with the prevalence of trimethoprim resistance in Enterobacteriaceae in England [5], while the use of trimethoprim in England was found to be associated with rates of urinary tract infection (UTI)-related bacteremia [7], with that association presumably mediated by treatment of trimethoprim-resistant UTIs with trimethoprim. Further instances of the association between the use of one antibiotic and prevalence of resistance to another antibiotic are shown in this paper (Tables 3 and 4). Those associations are likely related to patterns of cross-resistance for different pairs of antibiotics. For example, the high prevalence of cross resistance for fluoroquinolones and ES cephalosporins in *E. coli*-related UTIs described in the Introduction may help explain the association between the rates of fluoroquinolone prescribing and prevalence of resistance to ES cephalosporins in *E. coli* found in this paper (Table 4). Those mechanisms relating antibiotic use, resistance and severe outcomes support the utility of a granular evaluation of the relation between the rates of prescribing of different antibiotics, prevalence of antibiotic resistance and rates of severe outcomes associated with various syndromes with the aim of informing antibiotic prescribing guidelines.

The findings in this paper and other work (e.g. [4-10]) lead to the question whether antibiotic stewardship and replacement of certain antibiotics by others could bring about a reduction in the prevalence of resistance, as well as the rates of severe infections. Some evidence for that is provided by the reduction in fluoroquinolone and cephalosporin prescribing in the UK that was phased in starting in 2006. In particular, that reduction in prescribing was accompanied by the decrease in the prevalence of fluoroquinolone and cephalosporin resistance in certain Enterobacteriaceae in England [10], and prevalence of fluoroquinolone resistance in *C. difficile* [9], with a sharp reduction in the rates of *C. difficile* infection and deaths [9,33,34]. Use of fluoroquinolones was also found to be associated with MRSA acquisition [12-15], which may help explain the corresponding finding in our paper (Table 4). Rates of community-associated MRSA bacteremia in England dropped by more when half between 2006-2011 (Figures 1 and 3 in [33]), while the rates of community-associated MRSA invasive disease in the US were stable during the same period [35]. We also note that prevalence of fluoroquinolone (levofloxacin) non-susceptibility in community-associated MRSA isolates in the US in the recent years was high [36].

US FDA guidelines in recent years have recommended the restriction of fluoroquinolone use for certain conditions (such as uncomplicated UTIs), due to potential adverse effects [37]. The effect of those guidelines on fluoroquinolone prescribing, prevalence of resistance and the rates of associated severe bacterial infections remains to be seen. Additionally, those effects may depend on the choice of antibiotics used to replace fluoroquinolones, as there is high prevalence of resistance to other antibiotics in certain bacteria, as well as of cross-resistance between fluoroquinolones and other classes [16-19]. For example, replacing fluoroquinolones by penicillins in the US may not mitigate the rates of severe bacterial infections associated with certain pathogens, as suggested both by the US data and the UK experience. Levels of *E. coli* and *Klebsiella*-associated bacteremia continued to rise in England while the use of fluoroquinolones and cephalosporins was being reduced [10,33]. Rates of prescribing of amoxicillin-clavulanate (co-amoxiclav) in England rose significantly between 2006-2011 [38], and so did the rates of bacteremia associated with *E. coli* strains resistant to co-amoxiclav ([39,40]). In the US, prevalence of penicillin resistance in *E. coli*-associated urinary tract infections (UTIs) was found to be higher than the prevalence of fluoroquinolone resistance [16]. Antibiotic prescribing guidelines and practices that better account for patterns of resistance to different antibiotics in different bacteria are needed to mitigate the prevalence of antibiotic resistance and the rates of the associated severe bacterial infections. Along these lines, we note that nitrofurantoin is now generally recommended as the first-line option for the treatment of UTIs in the England [41], with prevalence of nitrofurantoin resistance in UTIs being lower in England compared to the prevalence of resistance to a number of other antimicrobials [40].

We found some negative associations between the rates of antibiotic prescribing and prevalence of antibiotic resistance. We note that other work found negative associations between the rates of nitrofurantoin, as well as macrolide prescribing and prevalence of resistance to trimethoprim in England [5]. These associations may partly reflect both model misspecification and the relation between the choice of an antibiotic and prevalence of antimicrobial resistance, with the use of some antibiotics rather than others contributing more to propagation of resistance to certain antibiotics (e.g. Table 2 in [16]). If the negative association between, for example, cephalosporin use and the prevalence of fluoroquinolone resistance in *P. aeruginosa* were purely attributable to the fact that cephalosporins are used instead of a drug that is more resistance-promoting and if our model were perfectly specified (i.e., the true relationship between the rates of antibiotic use and the prevalence of resistance were exactly additive on the logistic scale), then the multivariable model should account for this substitution effect and not estimate a negative coefficient for cephalosporin use. However, if (like all models) this model is somewhat mis-specified, the coefficient could be estimated as negative with this mechanism being operative. Other, more complex interactions are also possible explanations for the seemingly paradoxical negative associations; for example, one could envision that in states with high levels of cephalosporin use, the indigenous flora is more likely to be cephalosporin resistant and thus less susceptible to invasion or overgrowth of cephalosporin-resistant *P. aeruginosa* clones. While speculative, this general phenomenon is consistent with experimental evidence for other bacterial species [42]. At the same time, some of the negative associations between antibiotic use and resistance in our paper (particularly the high negative estimates for the relation between cephalosporin prescribing rates and prevalence of resistance to ES cephalosporins in *Klebsiella* spp., Tables 3 and 4) may also be affected by biases, including unmeasured confounding, though we also note the lack of univariate associations between prescribing rates for cephalosporins and prevalence of resistance to ES cephalosporins in *Klebsiella spp*. (Table 2), with negative point estimates for the correlations.

Our study has some additional limitations. The data in [26] do not contain information on several bacteria that are important sources of severe infections (e.g. methicillin-susceptible *S. aureus* and Streptococci), and on resistance to several types of antibiotics (e.g. macrolides and penicillins). Data on antibiotic prescribing in the whole population [25] were used in the regression model for which the outcomes were age-specific prevalence of resistance. We expect that this should generally introduce noise into the regression model, reducing precision rather than creating spurious associations. There is also some uncertainty in the relation between the prescribing of antibiotics in the outpatient setting, and the source of antimicrobial resistance in bacteria isolated in the CAUTI samples in hospitalized patients. While prevalence of resistance in bacteria in CAUTI samples is affected by the use of antibiotics and the prevalence of resistance in the community [24], it may also be affected by antibiotic administration in the hospital setting and practices related to urinary catheterization. While we tried to adjust for that by considering random effects for the US HSS regions, our results would benefit from independent analyses based on resistance data for bacterial specimens obtained in other settings. Such studies, including those involving individual-level data on antibiotic prescribing and resistance profiles for subsequent outcomes associated with bacterial infections should shed more light on the reasons behind the positive associations between antibiotic prescribing and antibiotic resistance found in this paper.

We believe that despite those limitations, our results give more support for the relation between antibiotic consumption and antibiotic resistance, at least for certain combinations of bacteria/antibiotics, highlighting the association between outpatient use of penicillins and fluoroquinolones in the US and prevalence of resistance to fluoroquinolones, as well as prevalence of bacterial organisms resistant to other antibiotics (e.g. ESBL-producing Enterobacteriaceae, and MRSA). Those results support the potential benefits of antibiotic stewardship, especially for penicillins and fluoroquinolones, and the need to improve the practices related to antibiotic, particularly penicillin and fluoroquinolone prescribing in the US, including tailoring antibiotic prescribing guidelines to patterns of resistance to different antibiotics.

